# Longitudinal white matter development in children is associated with puberty, attentional difficulties, and mental health

**DOI:** 10.1101/607671

**Authors:** Sila Genc, Charles B Malpas, Alisha Gulenc, Emma Sciberras, Daryl Efron, Timothy J Silk, Marc L Seal

## Abstract

The pubertal period involves dynamic white matter development. This period also corresponds with rapid gains in higher cognitive functions including attention, as well as increased risk of developing mental health difficulties. This longitudinal study comprised children aged 9-13 years (n=130). Diffusion magnetic resonance imaging (dMRI) data were acquired (b=2800 s/mm^2^, 60 directions) at two time-points. We derived measures of fibre density and morphology using the fixel-based analysis framework and performed a tract-based mixed-effects modelling analysis to understand patterns of white matter development with respect to age, pubertal stage, attentional difficulties, and internalising and externalising problems. We observed significant increases in apparent fibre density across a large number of white matter pathways, including major association and commissural pathways. We observed a linear relationship between fibre density and morphology with pubertal stage, in the right superior longitudinal fasciculus and in the right inferior longitudinal fasciculus. In terms of symptom severity, fibre density was positively associated with attentional dysfunction in the right uncinate fasciculus. Overall, white matter development across ages 9-13 years involved the expansion of major white matter fibre pathways, with key right-lateralised association pathways linked with pubertal development and attentional difficulties.

## 1. Introduction

Puberty is a critical period of development, marking the transition from childhood to reproductive maturity (Dorn et al. 2006). Brain structure is particularly sensitive to remodelling with exposure to pubertal hormones (Juraska and Willing 2017). Previous diffusion tensor imaging (DTI) studies have shown a strong link between advancing pubertal stage and greater white matter microstructural organisation (Ladouceur et al. 2012; Herting et al. 2012; Herting et al. 2017).

Evidence suggests that individual variability in pubertal timing is a strong predictor of mental health problems (Kaltiala-Heino et al. 2003). World-wide, the peak age of onset of psychiatric disorders is age 14 years (Kessler et al. 2005). The presence of internalising problems, including anxiety and depression, during this sensitive period of development pose risk for later case-level disorder (Shankman et al. 2009). Whether disrupted pubertal timing influences the onset of mental health problems and neurodevelopmental pathways, or vice versa, is not well understood. What is clear, however, is that these interrelated factors can alter the structural and functional reorganisation of the brain (Paus et al. 2008).

Alongside internalising problems, externalising disorders can co-exist with commonly occurring disorders in childhood, such as Attention-Deficit/Hyperactivity Disorder (ADHD) (Polanczyk et al. 2007). Whilst ADHD symptoms often present before pubertal onset, these symptoms can become more severe during the transition to adolescence, whereby increases in attentional difficulties and externalising symptoms can alter neuropsychological function (Langberg et al. 2008). Mapping these interrelated links between mental health and neurodevelopment is imperative. Previous DTI studies have shown a positive relationship between fractional anisotropy (FA) and ADHD symptom severity (Cooper et al. 2015) and a negative relationship between FA and internalising problems (Tromp et al. 2019; Hettema et al. 2012).

Recent advances in diffusion magnetic resonance imaging (dMRI) models facilitate the investigation of links between physical and behavioural phenotypes with brain development on a magnified neurobiological scale (Tamnes et al. 2017). The recently introduced fixel-based analysis (FBA) framework (Raffelt et al. 2017) and subsequent longitudinal modifications to this (Genc et al. 2018), allows the quantification of white matter fibre properties such as fibre density and morphology (see Box 1 for detailed interpretations). The primary measure of apparent fibre density (FD) reflects the total intra-axonal volume fraction per fibre pathway. The measure of fibre morphology encompasses both fibre cross-section (FC) and fibre-density and cross-section (FDC). FC is sensitive to macroscopic changes in the cross-sectional area perpendicular to a fibre bundle experienced during registration to a template image. FDC incorporates features of FD and FC, and acts as a surrogate marker of alterations to the capacity for information transfer across a fibre bundle.

#### Box 1: Interpretation of fixel-based metrics

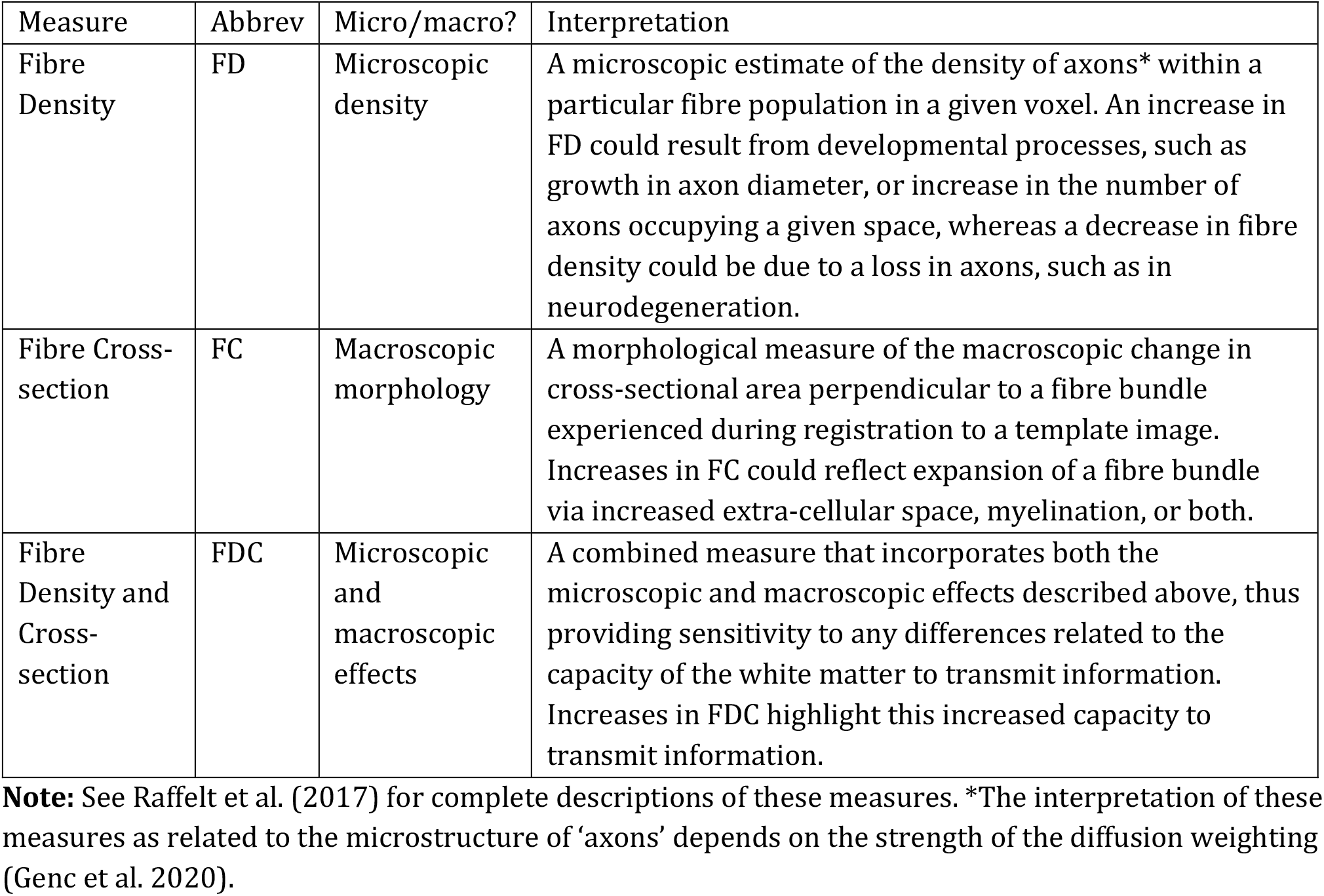

Compared to commonly used DTI metrics such as FA or mean diffusivity (MD), fixel-based FD is a stronger candidate for signifying microstructural organisation (Kelley et al. 2019), as it is both sensitive (at high b-values) to the intra-axonal signal (Genc et al. 2020), and specific to fibre populations (Raffelt et al. 2015). By comparison, FA measurements can be conflated by crossing fibres and extra-axonal signal contamination (Jones et al. 2013; Bach et al. 2014; Beaulieu 2009), making biophysical interpretations challenging.

With the use of robust neuroimaging studies using high-quality dMRI data, it is possible to unravel the specific neurobiological links between brain structure and developmental factors during this sensitive period of growth (Pines et al. 2020). To the best of our knowledge, this is the first longitudinal study to examine neurodevelopmental profiles of advanced dMRI measures with puberty, attention dysfunction, and mental health.

In this study, we investigate age, sex and puberty related development of white matter fibre density and morphology in commonly investigated white matter tracts. We then study the impact of symptom severity on white matter fibre properties. In order to understand these complex associations, we combine a tract-based approach with mixed-effects modelling upon a cohort of children with and without attentional difficulties. Rather than dichotomising groups at the extreme ends of pubertal stage, or mental health and attentional difficulties, we aimed to perform a dimensional analysis to shed light on how continuous patterns of physical and behavioural phenotypes relate to white matter fibre properties.

## 2. Methods

### Participants

This study reports on a sample of children aged 9-13 years recruited as part of the Neuroimaging of the Children’s Attention Project study (NICAP; see Silk et al. (2016) for a detailed protocol). This longitudinal study was approved by The Royal Children’s Hospital Melbourne Human Research Ethics Committee (HREC #34071). Briefly, children were initially recruited at 7 years of age as part of the Children’s Attention Project (CAP) study (Sciberras et al. 2013) from 43 socio-economically diverse primary schools distributed across the Melbourne metropolitan area, Victoria, Australia. Children underwent comprehensive assessment for ADHD at age 7 via the Diagnostic Interview Schedule for Children (DISC-IV) completed with parents face-to-face. Children were categorised as either meeting a negative or positive diagnosis for ADHD. A subsample of children from the CAP study, which either met or did not meet criteria for ADHD, were invited to participate in the neuroimaging study NICAP at age 10 (Figure S1).

At 10 years of age, children and their primary caregiver were invited for a 3.5-hour appointment at The Melbourne Children’s campus, which included a child assessment, parent questionnaire, mock scan, and MRI scan. Direct assessments and MRI scans were performed by a trained research assistant who was blind to the child’s diagnostic status. Children were invited for a follow-up appointment approximately 16 months following their initial visit (M = 16.14, SD = 2.37 months). Overall, only data from the two imaging time-points: time-point 1 (age: M = 10.4, SD = .44 years old) and time-point 2 (age: M = 11.7, SD = .51 years old), were included for analysis in the current study.

Written informed consent was obtained from the parent/guardian of all children enrolled in the study. Children were excluded from the study if they had a neurological disorder, intellectual disability, or serious medical condition (e.g. diabetes, kidney disease).

### Measures

The following measures were obtained at both imaging time-points. General intellectual ability was estimated using the Wechsler Abbreviated Scale of Intelligence (WASI) matrix reasoning sub-test (Wechsler 1999). The Connors 3 ADHD Index (10-items) was administered via parent survey (Conners et al. 2011), in order to capture the variation in ADHD symptom severity across time-points. The Strengths and Difficulties Questionnaire (SDQ) was administered in the form of a parent survey as a measure of emotional/behavioural difficulties (Goodman 1997). Using the responses from this questionnaire, the scores derived from the peer problems and emotional problems scales were added to generate a combined internalising difficulties score, and the scores from the hyperactivity and conduct scales were added to generate a combined externalising difficulties score (Goodman et al. 2010). The Pubertal Development Scale questionnaire (PDS; Petersen et al. (1988) was administered to parents, and a total PDS score(PDSS) combining features of adrenarche and gonadarche was computed for each imaging time-point (PDSS; Shirtcliff et al. (2009)). Additional information on the psychometric properties of these measures are summarised in Supplementary Information.

Child height and weight were measured using the average of two consecutive measurements to calculate a Body-Mass index (BMI) (kg/m^2^). Socio-economic status (SES) was determined using the Socio-Economic Indexes for Areas (SEIFA), based on Australian Census data.

### Image acquisition and pre-processing

Diffusion MRI data were acquired at two distinct time-points on a 3.0 T Siemens Tim Trio, at The Melbourne Children’s Campus, Parkville, Australia. Data were acquired using the following protocol: b = 2800 s/mm^2^, 60 directions, 4 volumes without diffusion weighting, 2.4 × 2.4 × 2.4 mm voxel size, echo-time / repetition time (TE/TR) = 110/3200 ms, multi-band acceleration factor of 3, acquisition matrix = 110 × 100, bandwidth = 1758 Hz. A total of 152 participants had longitudinal MRI data. Of those, 130 participants had useable diffusion MRI data, therefore the subsequent image processing and analysis was performed on these 130 participants with imaging data at two time-points (Figure S1).

All dMRI data were processed using MRtrix3 (v3.0RC3; Tournier et al. (2019)) using pre-processing steps from a recommended fixel-based analysis (FBA) pipeline (Raffelt et al. 2017). For each scan, these pre-processing steps were: denoising (Veraart et al. 2016); eddy, motion, and susceptibility induced distortion correction (Andersson and Sotiropoulos 2016); bias field correction (Tustison et al. 2010); and group-wise intensity normalisation. Data were then upsampled by a factor of 2, and a fibre-orientation distribution (FOD) was estimated in each voxel. Total intra-cranial volume for each T1-weighted image at each time-point was calculated using FreeSurfer (version 6) (Reuter et al. 2012). Images were visually inspected for motion artefact (assessed by the presence of Venetian blinding artefact), and whole datasets were excluded if excessive motion was present. In addition, we calculated mean frame-wise displacement using the FSL software library (v5.0.10) (Smith et al. 2004).

### Longitudinal template generation

In order to build an unbiased longitudinal template, we selected 40 individuals, with equal numbers of males and females, to first generate intra-subject templates. For each of these individuals, the time-point 1 and time-point 2 FOD maps were transformed to their midway space and subsequently averaged to generate an unbiased intra-subject template. The 40 intra-subject FOD templates were used as input for the population template generation step.

Following generation of the population template, each individual’s FOD image was registered to this longitudinal template (Raffelt et al. 2011), and the resulting transformed FOD within each template space voxel segmented to produce a set of discrete fixels. Reorientation of fixel directions due to spatial transformation, correspondence of these fixels with the template image, and derivation of FBA metrics, was performed as described previously (Genc et al. 2018). The output metrics fibre density (FD), fibre cross-section (FC), and fibre density and cross-section (FDC) (in template space) were then subjected to further statistical analysis.

### Tractography

We chose to delineate 7 key bilateral white matter fibre pathways, alongside 3 commissural bundles, which make up the John’s Hopkins University (JHU) white matter tractography atlas available in FSL. These pathways were delineated to segment fixels from the whole-brain fixel template which corresponded with our tracts of interest. To aid in the spatial identification of specific fibre bundles, we used a three-step process to improve specificity (Figure 1.1), summarised as follows:

- *Atlas registration:* The FA image derived from the JHU-ICBM atlas (Smith et al. 2004) was non-linearly transformed to our population template, and the resulting transformations were applied to warp the JHU tractography atlas to population template space.
- *Whole-brain tractogram overlay:* The whole-brain population-based tractogram was visualised in order to identify specific bundles. This tractogram was visualised as colour-coded directions, to further enable the identification of specific bundles (i.e. corticospinal tract runs inferior to superior, therefore is coloured blue).
- *Region of interest (ROI) selection:* We used a protocol defined in Wakana et al. (2007) to aid in the placement of inclusion ROIs for each major fibre bundle. We placed two separate inclusion ROIs in regions identified in the protocol, making sure they overlapped the JHU tractography atlas and whole brain tractogram. For bilateral tracts, the opposite hemisphere was used as an exclusion ROI. The whole brain tractography map was then edited using *tckedit*, and visualised, to check the anatomical correctness of the tract. In cases where spurious streamlines were present, these were rerun with an additional manually drawn exclusion ROI.

**Figure 1.1:**
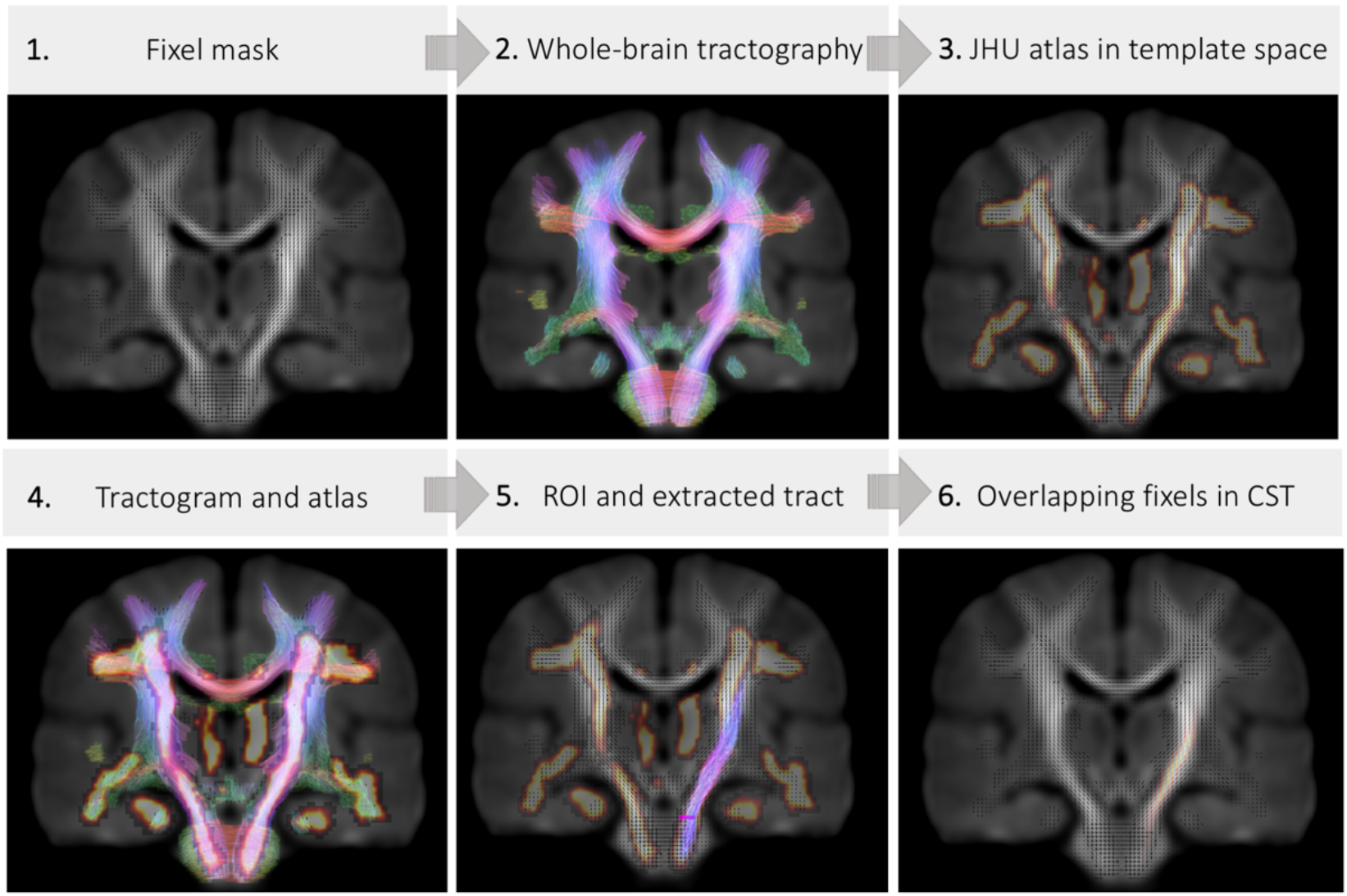
Protocol for defining fixels overlapping tracts of interest. Example shown is for a single ROI to delineate the left corticospinal tract

The subsequently generated 17 white matter tracts (Figure S2), also referred to as regions, were then converted to fixel maps using the whole-brain fixel template (*tck2fixel*), and binarised for statistical sampling of output maps (*mrthreshold*). For each participant at each time-point, we calculated mean FD, FC, and FDC values in each of the fixel masks derived from the 17 white matter tracts for subsequent statistical analyses. An example of these steps for one representative white matter tract can be visualised in Figure 1.1.

### Statistical analyses

All statistical analyses were performed within R (version 3.4.3). Data visualization was performed using *ggplot2* (Wickham 2016) and *raincloud* tools in R (van Langen 2020; M et al. 2019). To investigate age-related patterns of FBA metrics across the 17 tracts, and the effects of sex, puberty, and symptom severity, we applied linear mixed effects modelling using *lme4* (Bates et al. 2015). The most parsimonious model was selected based on lowest Akaike Information Criterion (AIC) values.

First, we tested the interaction between FBA metrics across the 17 white matter tracts with age and sex. Then, for each white matter region we tested whether age, sex, and puberty predicted FD, FC and FDC values. We repeated the same analysis for age, sex, and symptom severity (internalising symptoms, externalising symptoms, ADHD symptoms). Finally, we tested whether the longitudinal change in pubertal stage predicted the longitudinal change in FBA measures. Full details of each model are summarised in Table S1.

We computed bootstrapped 95% confidence intervals (n=5000 simulations) and report these for each mixed-model coefficient as β [95% CI]. Evidence for an association is represented when confidence intervals do not cross zero. We additionally report p-values, adjusted using False Discovery Rate (FDR) correction (*p_FDR_*; Benjamini and Hochberg (1995)).

## 3. Results

### Participant characteristics

Differences in participant characteristics over the 16-month follow-up period are reported in Table 1.1 and visualised in Figure 1.2. We observed developmental increases in physical characteristics such as BMI and PDSS (p < .001), and in WASI matrix reasoning raw score (p < .001). Behaviourally, we observed an increase in internalising symptoms (p < .001) but no change in externalising symptoms (p > .05) assessed by the SDQ. There was some evidence for a decrease in ADHD symptoms over time (p = .04).

**Figure 1.2:**
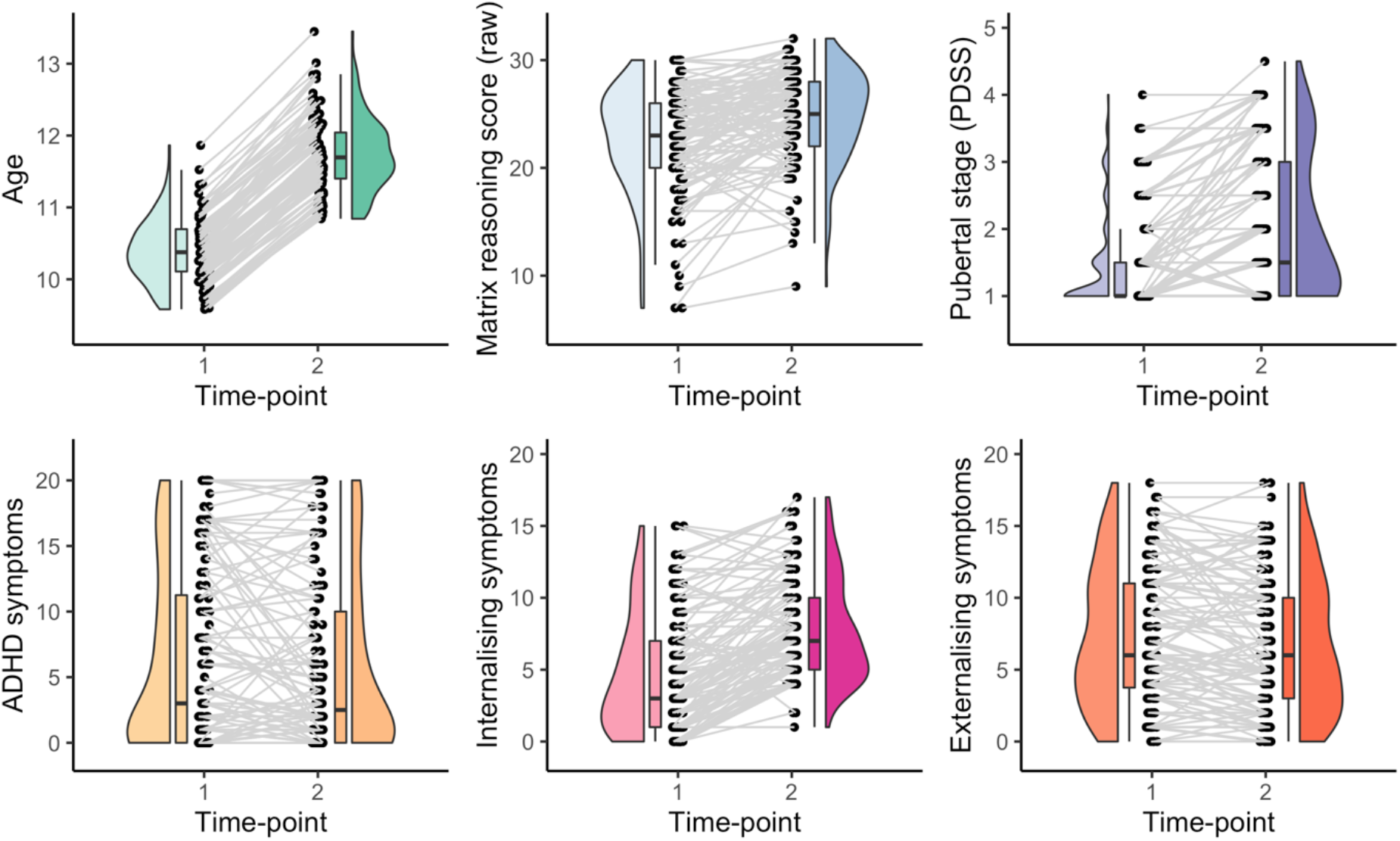
Change in participant characteristics over the 16-month follow-up period. Longitudinal data points are connected by a line.

**Table 1.1:** Change in participant characteristics over the follow-up interval.

| Variable | Time-point 1<br>M(SD) |  | Time-point 2<br>M(SD) |  | Difference<br>*p-value* |
| --- | --- | --- | --- | --- | --- |
|  | Female | Male | Female | Male |  |
| Age, years | 10.36 (.40) | 10.39 (0.46) | 11.67 (.44) | 11.75 (.55) | < .001 |
| Socio-economic status (SES) | 1007 (46) | 1022 (44) | 1007 (46) | 1021 (44) | .32 |
| Body Mass Index, kg/m<sup>2</sup> | 20 (3.5) | 19 (3.8) | 21 (4.0) | 20 (4.3) | < .001 |
| Pubertal stage, PDSS | 2.0 (.9) | 1.2 (.4) | 3 (.9) | 1.5 (.7) | < .001 |
| SDQ internalising | 4.8 (4) | 4.5 (4) | 8.1 (4) | 7.4 (3) | < .001 |
| SDQ externalising | 5.7 (5) | 7.7 (5) | 5.1 (4) | 7.4 (5) | .19 |
| ADHD symptoms | 3.8 (6) | 7.5 (7) | 3.3 (5) | 6.9 (7) | .04 |
| Matrix reasoning, raw score | 22 (4.7) | 23 (4.9) | 24 (3.4) | 25(4.6) | < .001 |

Difference between time-points calculated using a paired samples t-test. N(females) = 47; N(males) = 83. M = mean, SD = standard deviation

We assessed whether there was evidence for regional differences in age-related patterns of white matter microstructure (Figure 1.3), and associations between our phenotypic variables of interest. We observed a significant region by age interaction, suggesting that region-specific age-related patterns exist. To investigate these region-specific relationships, we computed multiple linear mixed-effects models for each white matter tract. Results from the mixed-effects modelling analysis are presented for each predictor (Tables 1.2 & 1.3; Figures 1.3 & 1.4).

**Figure 1.3:**
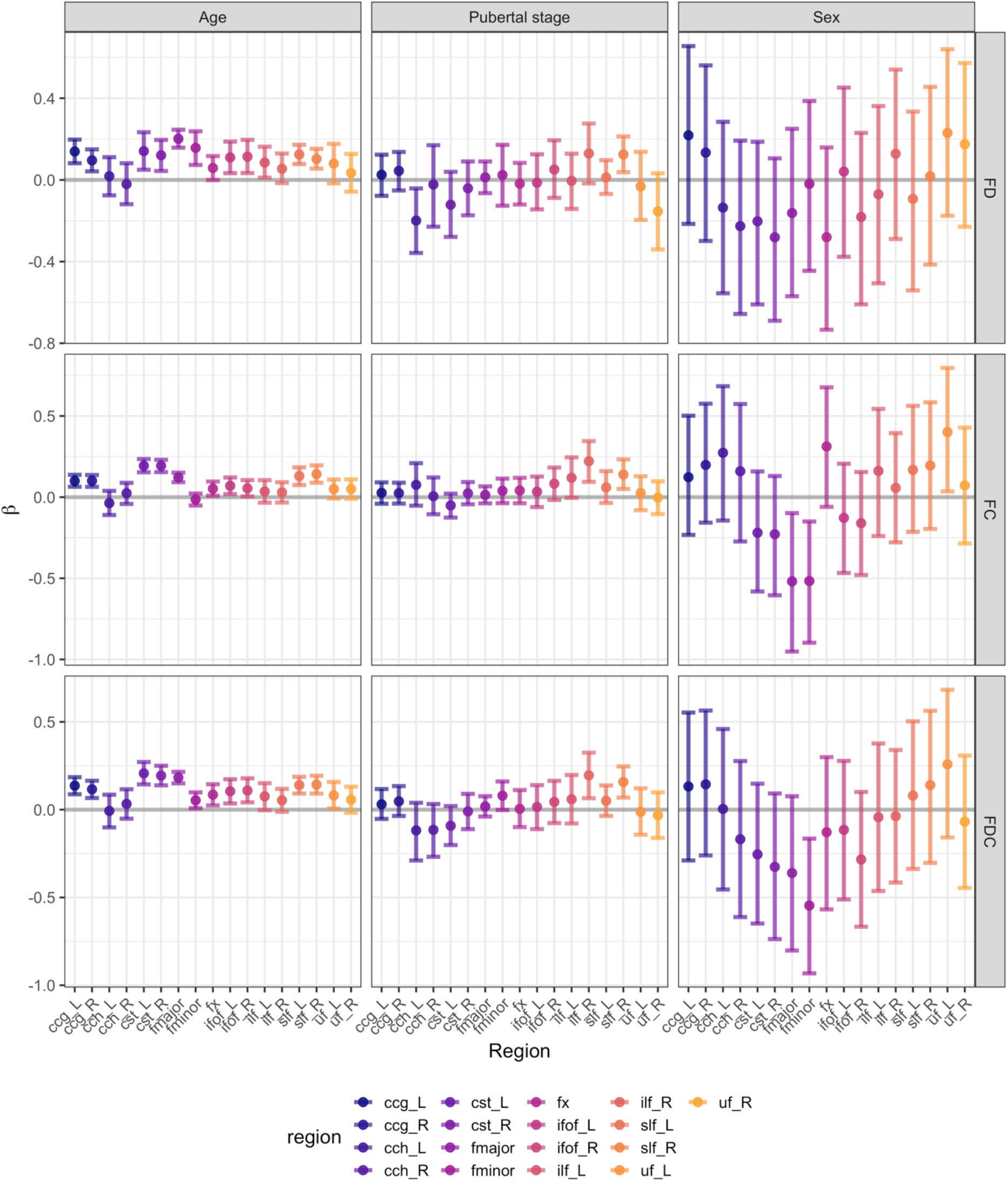
Relationships between participant characteristics and fibre microstructure across all white matter tracts. 95% confidence intervals which do not cross zero suggest a relationship between that variable and the metric of interest.

**Table 1.2:**
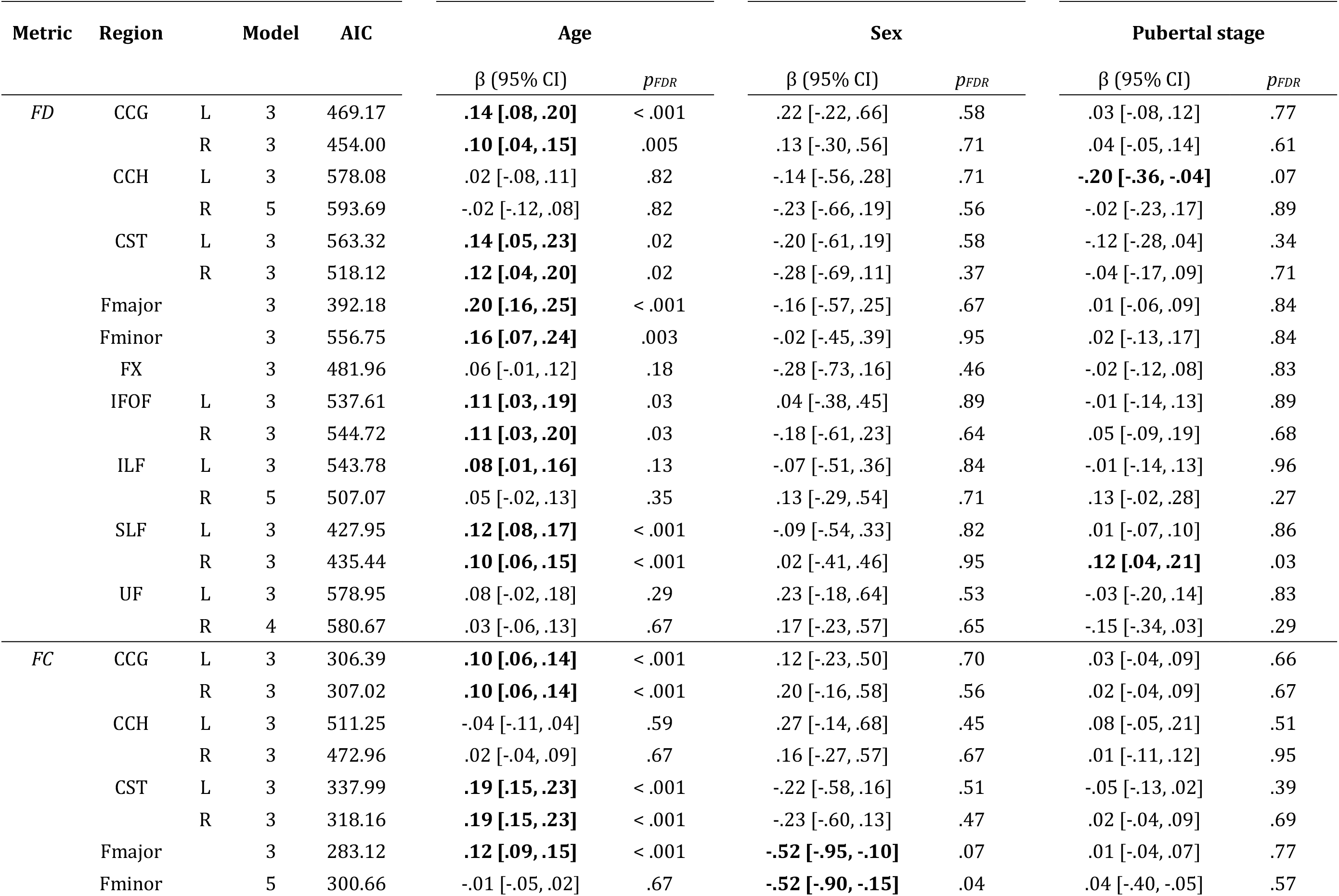
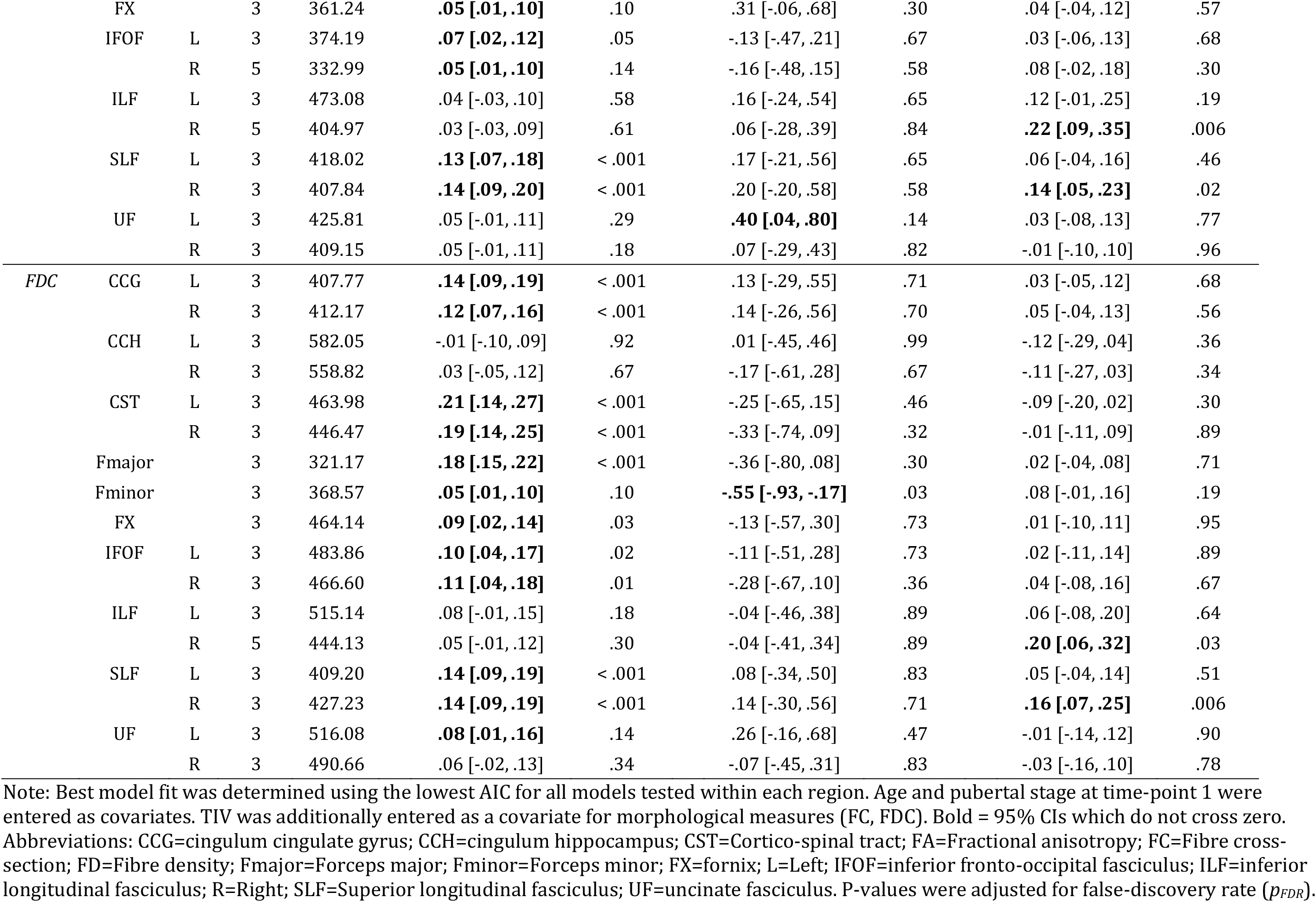
Relationship between white matter microstructure with age, sex, and pubertal stage.

**Table 1.3:**
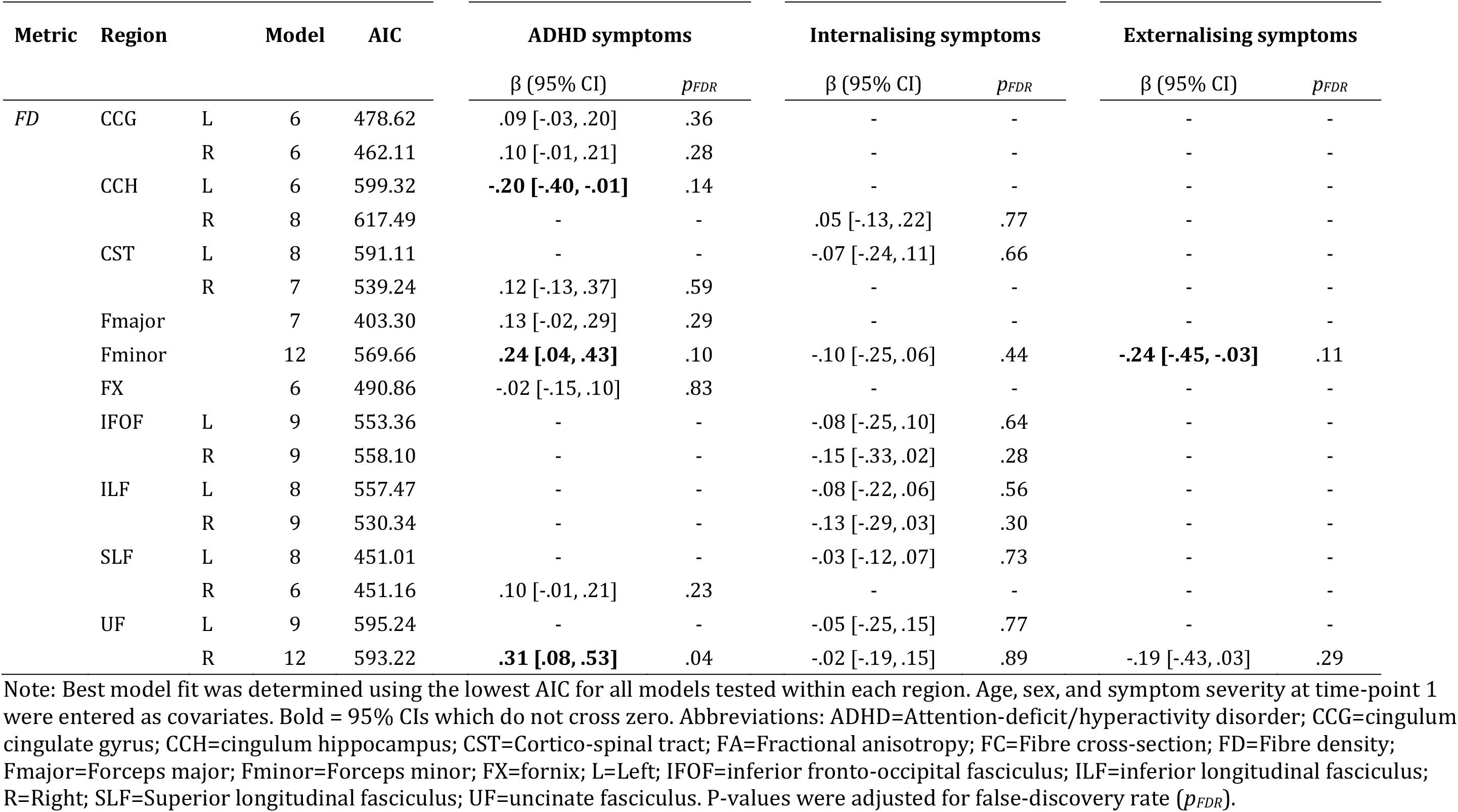
Relationship between white matter microstructure and symptom severity.

| Metric | Region |  | Model | AIC | ADHD symptoms |  | Internalising symptoms |  | Externalising symptoms |  |
| --- | --- | --- | --- | --- | --- | --- | --- | --- | --- | --- |
| | | | | | $\beta$ (95% CI) | $p_{FDR}$ | $\beta$ (95% CI) | $p_{FDR}$ | $\beta$ (95% CI) | $p_{FDR}$ |
| FD | CCG | L | 6 | 478.62 | .09 [-.03, .20] | .36 | - | - | - | - |
|  |  | R | 6 | 462.11 | .10 [-.01, .21] | .28 | - | - | - | - |
|  | CCH | L | 6 | 599.32 | **-.20 [-.40, -.01]** | .14 | - | - | - | - |
|  |  | R | 8 | 617.49 | - | - | .05 [-.13, .22] | .77 | - | - |
|  | CST | L | 8 | 591.11 | - | - | -.07 [-.24, .11] | .66 | - | - |
|  |  | R | 7 | 539.24 | .12 [-.13, .37] | .59 | - | - | - | - |
|  | Fmajor |  | 7 | 403.30 | .13 [-.02, .29] | .29 | - | - | - | - |
|  | Fminor |  | 12 | 569.66 | **.24 [.04, .43]** | .10 | -.10 [-.25, .06] | .44 | **-.24 [-.45, -.03]** | .11 |
|  | FX |  | 6 | 490.86 | -.02 [-.15, .10] | .83 | - | - | - | - |
|  | IFOF | L | 9 | 553.36 | - | - | -.08 [-.25, .10] | .64 | - | - |
|  |  | R | 9 | 558.10 | - | - | -.15 [-.33, .02] | .28 | - | - |
|  | ILF | L | 8 | 557.47 | - | - | -.08 [-.22, .06] | .56 | - | - |
|  |  | R | 9 | 530.34 | - | - | -.13 [-.29, .03] | .30 | - | - |
|  | SLF | L | 8 | 451.01 | - | - | -.03 [-.12, .07] | .73 | - | - |
|  |  | R | 6 | 451.16 | .10 [-.01, .21] | .23 | - | - | - | - |
|  | UF | L | 9 | 595.24 | - | - | -.05 [-.25, .15] | .77 | - | - |
|  |  | R | 12 | 593.22 | **.31 [.08, .53]** | .04 | -.02 [-.19, .15] | .89 | -.19 [-.43, .03] | .29 |

Note: Best model fit was determined using the lowest AIC for all models tested within each region. Age, sex, and symptom severity at time-point 1 were entered as covariates. Bold = 95% CIs which do not cross zero. Abbreviations: ADHD=Attention-deficit/hyperactivity disorder; CCG=cingulum cingulate gyrus; CCH=cingulum hippocampus; CST=Cortico-spinal tract; FA=Fractional anisotropy; FC=Fibre cross-section; FD=Fibre density; Fmajor=Forceps major; Fminor=Forceps minor; FX=fornix; L=Left; IFOF=inferior fronto-occipital fasciculus; ILF=inferior longitudinal fasciculus; R=Right; SLF=Superior longitudinal fasciculus; UF=uncinate fasciculus. P-values were adjusted for false-discovery rate ( $p_{FDR}$ ).

### Age, sex, and pubertal stage

Age related increases in fibre density (FD) over time were localised to the bilateral CCG, bilateral CST, forceps major and minor, bilateral IFOF, left ILF, and bilateral SLF. Significant increases in fibre cross-section (FC) were observed in: bilateral CCG, bilateral CST, forceps major, fornix, bilateral IFOF, and bilateral SLF. Increases in fibre density and cross-section (FDC) were observed in bilateral CCG, bilateral CST, forceps major and minor, fornix, bilateral IFOF, bilateral SLF, and left UF. All regions listed significantly increased in fibre density and morphology over time, and no regions decreased. The longitudinal changes in common fibre pathways changing over time are visualised in Figure 1.4.

**Figure 1.4:**
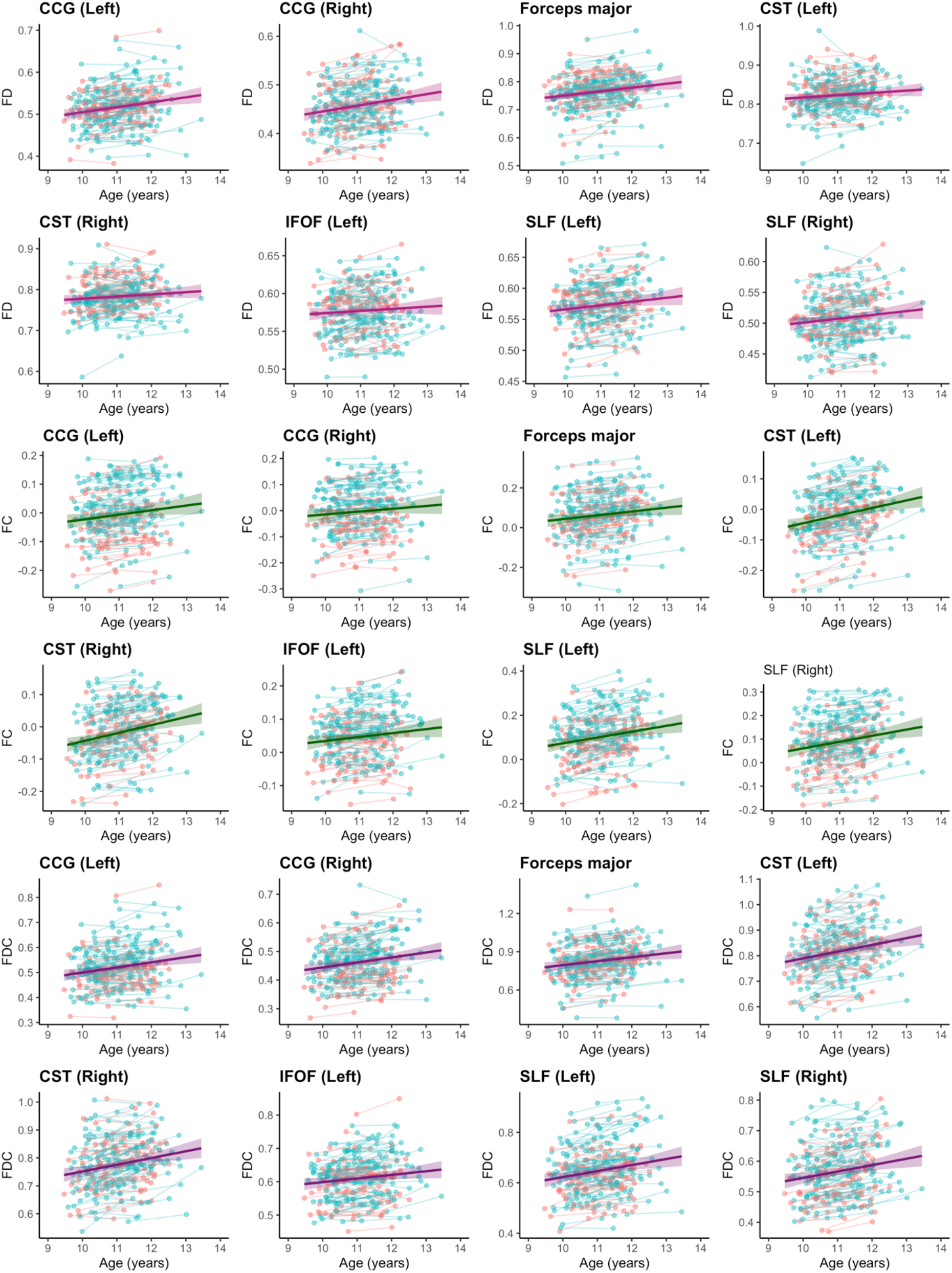
Longitudinal change in white matter fibre properties for regions with significant age-related increases over time. Blue = boys, Red = girls

We observed no significant sex differences in fibre density across any of the regions studied. For fibre cross-section, females had higher FC than males in the forceps major, β [95% CI] = −.52 [−.95, −.10], forceps minor, β [95% CI] = −.52 [−.90, −.15], whereas males had higher FC than females in the left uncinate fasciculus, β [95% CI] = .40 [.04, .80]. In addition, females had higher FDC in the forceps minor compared with males, β [95% CI] = −.55 [−.93, −.17].

We observed a positive relationship between pubertal stage and fibre properties in the right superior longitudinal fasciculus for FD, β [95% CI] = .12 [.01, .21], FC, β [95% CI] = .14 [.05,.23], and FDC, β [95% CI] = .16 [.07, .25], and for the right inferior longitudinal fasciculus for FC, β [95% CI] = .22 [.09, .35], and FDC, β [95% CI] = .20 [.06, .32]. We observed a negative relationship between pubertal stage and FD in the left cingulum hippocampus, β [95% CI] = −.20 [−.36, −.04].

Total intra-cranial volume (TIV) was a significant predictor of: FD for the forceps minor, right ILF, and right UF; FC over all of the white matter tracts studied; and FDC for all regions apart from the left CCH.

For a select number of regions, models including age and/or sex interaction terms were preferred (e.g. right CCH, right IFOF, right ILF, right UF; Table 1.2). We observed an interaction between sex and pubertal stage on fibre density in the right uncinate fasciculus, β [95% CI] = .25 [.01, .48]. We observed an age * PDSS interaction in the right inferior longitudinal fasciculus for FC and FDC, β [95% CI] = −.08 [−.14, −.03].

### Symptom severity

The relationship between symptom severity and white matter microstructure is summarised in Table 1.3. The main findings are represented in Figure 1.5. We found evidence for a positive association between FD and ADHD symptoms in the forceps minor, β [95% CI] = .24 [.04, .43], and the right UF, β [95% CI] = .31 [.08, .53], and a negative association in the left cingulum hippocampus, β [95% CI] = −.20 [−.40, −.01]. We observed a negative association between externalising symptoms and fibre density in the forceps minor, β [95% CI] = −.24 [−.45, −.03]. We observed no significant relationship between fibre morphology and symptom severity for any of the tracts studied. For a select number of regions, models including age and/or sex interaction terms were preferred (e.g. right CST, forceps major and minor, bilateral IFOF, right ILF, bilateral UF). We observed an interaction between sex and internalising symptoms on fibre density in the right inferior fronto-occipital fasciculus, β [95% CI] = .21 [.03, .39].

**Figure 1.5:**
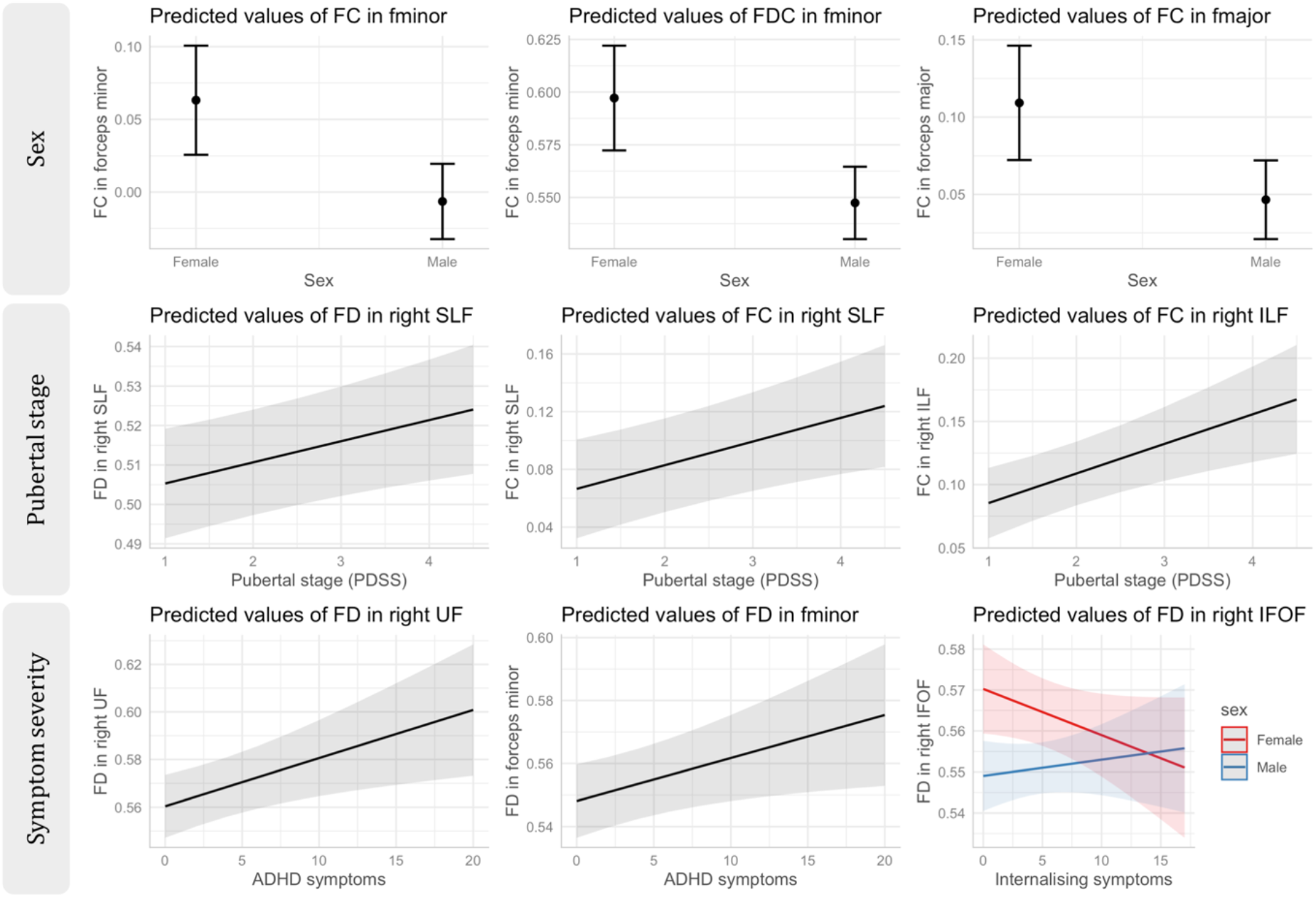
Marginal effects plots for the main mixed-model results for sex, pubertal stage, and symptom severity.

### Pubertal progression

We observed an interaction between the change in pubertal stage and age, with the longitudinal change in fibre density over time for: the left corticospinal tract, β [95% CI] = .19 [.04, .35], left inferior fronto-occipital fasciculus, β [95% CI] = .25 [.07, .42], left inferior longitudinal fasciculus, β [95% CI] = .19 [.01, .37], and the left uncinate fasciculus, β [95% CI] = .24 [.06, .42]. For these regions, a larger change in PDSS score combined with older age resulted in a larger increase in fibre density.

### Impact of motion

Analyses were rerun with frame-wise displacement included to mitigate any impact of motion on our findings, particularly given the nature of the sample and potential for greater motion artefact in children with attention/hyperactivity phenotypes. The results for the linear mixed-effects models remained unchanged (Table S2), and we observed no main effect of motion on the models tested for fibre density in each tract. We did observe an effect of motion on the morphological measures (FC and FDC) for some tracts, suggesting that motion artefact can influence the computation of morphological measures, however, this had no bearing on the final results.

## 4. Discussion

In our longitudinal diffusion MRI study of 130 children across two time-points, we identify regional patterns of white matter fibre density and morphology development related to pubertal stage, attentional difficulties, and mental health. These findings open up new avenues for investigating the interaction between developmental factors, to unravel multiple contributions to altered neurodevelopmental pathways.

### Developmental patterns

We observed age-related development of fibre density in the bilateral cingulum cingulate gyrus, bilateral corticospinal tract, forceps major and minor, bilateral inferior fronto-occipital fasciculus, and bilateral superior longitudinal fasciculus. Similar patterns were observed for fibre cross-section and fibre density and cross-section. The region-specific development we observed is consistent with findings from many diffusion tensor imaging studies (see Tamnes et al. (2018) for a full review) and fixel-based analyses (Genc et al. 2018) in typically developing children of this age range. This may suggest that regional fibre development is consistent in this age range (9 - 13 years), regardless of variations due to pubertal stage, attentional difficulties, and mental health. Regions such as the bilateral cingulum hippocampus, right inferior longitudinal fasciculus, and right uncinate fasciculus may have delayed maturation, completing maturation closer to the completion of adolescence and early adulthood (Lebel and Beaulieu 2011).

In terms of sex differences in microstructure, a number of longitudinal developmental studies have also reported no sex differences in white matter development using DTI (Brouwer et al., 2012; Krogsrud et al., 2016; Lebel and Beaulieu, 2011; Tamnes et al., 2010), consistent with our findings. However, some studies of older samples studies have shown that boys have higher FA than girls in a number of associative white matter pathways (Seunarine et al. 2016; Schmithorst et al. 2008; Herting et al. 2012). Given our analyses are focused on data acquired at high diffusion weightings sensitised to the intra-axonal signal, discrepancies in findings could be due to differences in sensitivity to various neurobiological properties, for example myelination and extracellular properties.

The sex differences in fibre morphology observed in the forceps minor suggest that females have greater capacity for information transfer across this white matter tract. These differences could plausibly be a signature of anatomical sex differences between males and females, potentially induced by early perinatal exposure to testosterone (Sisk and Foster 2004). Neuroimaging studies have shown that testosterone and oestradiol impact brain volume across adolescence (Herting et al. 2014). Additionally (or alternatively) these differences could be perpetuated by the onset of pubertal processes and rising hormone levels in females, compared with males. Oestrogen receptors exist on oligodendrocytes, which may provide an avenue for myelin production (Zhang et al. 2004). Whilst we have not directly studied myelination in this study, this may be one explanation for the sex differences in fibre morphology observed in anterior and posterior callosal tracts.

### Pubertal stage and progression

A positive association between pubertal stage and both fibre density and morphology was observed in the right superior longitudinal fasciculus. This suggests a linear relationship between pubertal development and fibre maturation, likely driven by the expansion of fibre bundles by virtue of axonal growth and myelination. The superior longitudinal fasciculus is a white matter fibre most associated with language ability, semantic memory, and executive function, in developmental and adult populations. It anatomically connects two regions important for language - Broca’s and Wernicke’s area, and the relationship between language ability and microstructure in the SLF has been extensively reported (Catani et al. 2005). Similarly, we observed that fibre morphology of the right inferior longitudinal fasciculus exhibited a positive relationship with pubertal stage.

The pubertal period involves regulation and remodelling of white matter (Herting et al. 2017; Herting et al. 2012) with rising adrenal and gonadal (Maninger et al. 2009; Perrin et al. 2008; Pesaresi et al. 2015; Pangelinan et al. 2016) hormones. Previous studies have shown puberty-related increases in FA in the right insular gyrus with physical development (Herting et al. 2012) and decreases in MD in the superior and inferior longitudinal fasciculi (Menzies et al. 2015). This is consistent with our findings of a positive relationship between pubertal stage and fibre density and morphology in the right SLF, and fibre morphology in the right ILF. MD is sensitive to isotropic diffusion in the extracellular space, therefore any increases in intra-axonal signal fraction (FD) can be reflected by decreases in MD, assuming myelin thickness does not change drastically.

A recent longitudinal study of non-human primate (marmoset) brain development investigated grey matter and white matter changes over 3 - 7 time-points across the pre-pubertal, pubertal, and post-pubertal periods (Sawiak et al. 2018). Their results showed that the most dynamically developing white matter tracts in the early-pubertal period are the splenium of the corpus callosum, and ILF. These results are in line with our previous cross-sectional results, whereby fibre density appears to be increasing in the splenium in response to pubertal onset (Genc et al. 2017), and with the current results as we observed increasing fibre morphology in the right ILF with pubertal stage. During the pubertal period, regions experiencing most marked white matter tract thickening were localised to the SLF (Sawiak et al. 2018), consistent with evidence that the SLF does not reach full maturation until the post-pubertal stage (Ladouceur et al.2012). Overall, we have replicable evidence that the splenium of the corpus callosum and right inferior longitudinal fasciculus are sensitive to pubertal timing, and the right superior longitudinal fasciculus is sensitive to pubertal progression. These findings additionally highlight the sensitivity of advanced white matter fibre metrics such as fibre density, as these metrics appear to be sensitive to region-specific puberty-mediated white matter changes.

In terms of pubertal progression, it appeared that older children with a larger change in PDSS score resulted in larger increases in fibre density in the left corticospinal tract, left inferior fronto-occipital fasciculus, left inferior longitudinal fasciculus, and left uncinate fasciculus. This may suggest that the left-lateralised association tracts (e.g. IFOF, ILF, UF) with known protracted development in late adolescence (Von Der Heide et al. 2013) are developing earlier due to faster pubertal progression.

### Symptom severity

Greater attentional difficulties were associated with higher fibre density in the right uncinate fasciculus and forceps minor. Interpreting these findings in the wider context of previous findings is difficult as the ADHD literature surrounding macro-or micro-structural development in this population is consistently inconsistent (van Ewijk et al. 2012). Some studies report that microstructural organisation is greater in ADHD cases compared with controls (Peterson et al. 2011; Silk et al. 2009), whereas other studies report the exact opposite (Ashtari et al. 2005; Nagel et al. 2011; Wu et al. 2019). The underlying factors could be neurobiological, whereby children with attentional difficulties have altered patterns of microstructure, or these differences could also be induced by limitations of analysis techniques (Bach et al. 2014) or motion-artefact causing local increase/decrease in FA (Aoki et al. 2018). We have attempted to mitigate for motion artefact with the use of frame-wise displacement as a covariate, which did not appear to influence the main results. However, more robust measures such as temporal signal-to-noise ratio (TSNR) may be more appropriate descriptors of motion artefact (Roalf et al. 2016).

In the context of the current study, our findings are concordant with previous reports of higher FA with greater ADHD symptom severity (Cooper et al. 2015), with an added level of confidence of a longitudinal study design. Given the transient nature of ADHD symptomatology, where children can transition in and out of diagnostic categories, longitudinal studies are imperative to tease apart robust relationships between symptomatology and microstructure - not just static associations.

The observed sex interaction with internalising symptoms on fibre density in the right inferior fronto-occipital fasciculus suggests that females have a stronger negative relationship between internalising symptoms and fibre density compared with males. Similar findings using DTI have been reported in the context of anxiety disorders (Hettema et al. 2012; Phan et al. 2009; Tromp et al. 2019). Females could potentially be at more risk for interrupted fibre development due to the earlier influx of adrenal and gonadal hormones, resulting in increased internalising difficulties (Whittle et al. 2015) and impacting region-specific timing of neurodevelopment (Neufang et al. 2009; Ahmed et al. 2008).

### Relationship with brain volume

Fibre cross-section is inherently a morphological measure of the macroscopic change in cross-sectional area perpendicular to a fibre bundle experienced during registration to a template image (Raffelt et al. 2017). This finding is of special interest to users of this analysis framework and derived metrics, as any variable associated with ICV can potentially conflate interpretations on changes in fibre cross-section (Smith et al. 2019). For example, it is known that on average (for a given age), males have greater ICV than females. Without adjusting for ICV in fibre cross-section estimates, sex differences could be attributed to biologically relevant differences, when in fact the differences may purely due to large-scale anatomical differences in head size (Luders et al. 2014). Similar patterns are observed in the analyses of fibre density and cross-section, given that this metric is calculated from fibre cross-section. Therefore, the main message of these results, are that these metrics must be interpreted with caution, and that morphological measures should be adjusted appropriately for ICV.

### Limitations and future directions

Operating within the recommended fixel-based analysis framework may not necessarily be optimal for the tract segmentation method described in the current study. The fixel mask generation step is optimised to ensure the core or ‘deep’ white matter is included for subsequent fixel segmentation and reducing the number of multiple comparisons. However, in the context of a tract-based analysis, these steps may result in a smaller area of tract being delineated, and therefore we may potentially miss subtle variations of fibre density and morphology in the more peripheral aspects of a tract. Future studies might consider adopting a looser threshold for fixel segmentation to capture peripheral white matter more effectively.

Our measures of symptom severity may be biased as our sample inherently includes children which have either met or not met criteria for ADHD three years prior to neuroimaging. Therefore, although our community-based sample captures a wide range of symptom severity for attentional difficulties, and internalising and externalising symptoms, we may be missing out on high-risk children which fit ‘in-between’ these two ends of the spectrum of symptom severity. Whilst we have attempted to covary for symptom severity in a dimensional fashion to account for individual variation within groups, future studies should perform a larger population-level analysis to account for a wider range in attentional difficulties.

Our approach studied separate white matter pathways to determine the best fitting model for each region. Whilst useful for ensuring each region is studied with the appropriate predictors, the problem of multiple comparisons can influence the interpretability of results. Dimensionality reduction methods such as principal components analysis (Chamberland et al. 2019) or canonical correlation analysis (Ball et al. 2018) has proven useful in these scenarios, in order to capture components of microstructural attributes which vary together over development.

### Conclusion

We summarise our longitudinal findings with four important conclusions: (1) white matter development over the ages of 9 – 13 involves dynamic increases in fibre density and morphology in a large range of developmentally-sensitive white matter fibre pathways, (2) pubertal stage is positively associated with fibre density and morphology in the right superior longitudinal fasciculus, (3) intra-cranial volume is highly predictive of fibre morphology and should be modelled appropriately, and (4) attentional dysfunction is associated with fibre density in the uncinate fasciculus. These results shed light on key fibre developmental properties with age and puberty, and how these may relate to attention dysfunction and mental health.

## Abbreviations

ADHD: Attention-deficit/hyperactivity disorder
CCG: cingulum cingulate gyrus
CCH: cingulum hippocampus
CSD: Constrained Spherical Deconvolution
CST: Cortico-spinal tract
FA: Fractional anisotropy
FBA: Fixel-based analysis
FC: Fibre cross-section
FD: Fibre density
FDC: Fibre density and cross-section
FOD: Fibre orientation distribution
Fmajor: Forceps major
Fminor: Forceps minor
Fx: fornix
IFOF: inferior fronto-occipital fasciculus
ILF: inferior longitudinal fasciculus
MRI: Magnetic resonance imaging
NICAP: Neuroimaging of the Children’s Attention Project
PDS: Pubertal development scale
SEIFA: Socio-Economic Indexes for Areas
SES: Socio-economic Status
SLF: Superior longitudinal fasciculus
UF: uncinate fasciculus

## 5. Declaration

All authors declare no real or potential conflicts of interest.

## 6. Acknowledgements

Data used in the preparation of this article were obtained from the NICAP study (National Health and Medical Research Council; project grant #1065895). This research and analysis was conducted within the Developmental Imaging research group, Murdoch Children’s Research Institute, supported by the Victorian Government's Operational Infrastructure Support Program, and The Royal Children’s Hospital Foundation devoted to raising funds for research at The Royal Children’s Hospital. SG was supported by an Australian Government Research Training Program (RTP) scholarship. ES was supported by an NHMRC Career Development Award (1110688).

We extend our gratitude to the families that have participated for a number of years in this longitudinal study. We thank The Royal Children’s Hospital Medical Imaging staff for their assistance and expertise in the collection of the MRI data included in this study.

